# Most inhibitors with defined yeast targets have limited activity against the microsporidian *Nematocida parisii*

**DOI:** 10.64898/2026.06.09.731163

**Authors:** Qingyuan Huang, Guoqing Pan, Jie Chen, Aaron W. Reinke

## Abstract

Microsporidia are obligate intracellular parasites that infect diverse animals. Budding yeast has long been used to define core molecular pathways and inhibitors that target them. To test whether these compounds can target conserved pathways in the microsporidian *Nematocida parisii*, we assessed 15 inhibitors with defined yeast targets that had not previously been tested for effects on infection of *Caenorhabditis elegans*. Most showed little activity against *N. parisii*, except tunicamycin. These results identify tunicamycin as a potential tool for studying endoplasmic reticulum stress in microsporidia, while indicating that most yeast-targeted inhibitors are largely ineffective in this system.

## 1. Introduction

Microsporidia are a large phylum of obligate intracellular parasites that infect humans and agriculturally important animals such as honeybees, silkworms, and shrimp (Wadi and Reinke, 2020). At least 17 species are known to infect humans, causing symptoms such as diarrhea and, in some cases, lethal infections, while infections of silkworms and shrimp have caused substantial economic losses (Franzen, 2008; Han et al., 2021; Shinn et al., 2018). Because of the threat microsporidia pose to health and food security, there is a need for effective inhibitors, yet the molecular targets of most candidate compounds remain unknown (Huang et al., 2026b). In contrast, the budding yeast *Saccharomyces cerevisiae* has small-molecule inhibitors with well-defined protein targets that have been used extensively to manipulate and dissect core cellular pathways (Ottilie et al., 2022). Many of these pathways are conserved across eukaryotes, raising the possibility that such inhibitors could be repurposed to probe the corresponding pathways in microsporidia (Parsons et al., 2004). The nematode *Caenorhabditis elegans* and its natural microsporidian parasite *Nematocida parisii* have been developed as a convenient whole-animal model for identifying microsporidia inhibitors, characterizing their effects, and determining protein targets (Huang et al., 2026b, 2025; Murareanu et al., 2022). Infection by *N. parisii* suppresses *C. elegans* progeny production, providing an easily scored phenotype, and over 10,000 compounds have been screened in this system, identifying inhibitors that inactivate spores, block germination, or prevent proliferation (Huang et al., 2026a). However, whether inhibitors with characterized targets in yeast are active against *N. parisii*, and whether activity can be predicted from the conservation of the target, has not been examined.

To address this, we tested a panel of small-molecule inhibitors with defined yeast targets for their ability to inhibit *N. parisii* infection of *C. elegans*. We assembled 18 inhibitors, annotated the *N. parisii* ortholog of each yeast target along with its degree of conservation, and measured the effect of each compound on infection and host gravidity across a range of concentrations. Most inhibitors showed weak anti-microsporidia activity. One compound, tunicamycin, strongly inhibited *N. parisii* infection, suggesting it may be a useful tool for studying endoplasmic reticulum stress during microsporidia infection.

## 2. Materials and Methods

### *C. elegans* maintenance and *E. coli* culture

The wild-type *C. elegans* strain N2 was maintained on nematode growth medium (NGM) plates seeded with *Escherichia coli* OP50 as a food source (Tamim El Jarkass et al., 2022). To generate age-synchronized populations, gravid adults were washed from plates and dissolved in a sodium hypochlorite/sodium hydroxide solution, and the released embryos were allowed to hatch in M9 buffer at 21°C for 18–24 h. To prepare OP50, a frozen stock was streaked onto LB agar and grown overnight at 37°C, after which a single colony was used to inoculate a liquid LB culture that was grown for 18 h at 37°C with shaking. This culture was concentrated tenfold by centrifugation and stored at 4°C.

### Chemicals

All compounds were obtained from Cayman Chemical, with the exception of dexrazoxane and albendazole, which were obtained from Sigma-Aldrich. Stock solutions were prepared in DMSO and stored at −80°C.

### *N. parisii* spore preparation

*N. parisii* spores were prepared as described previously (Huang et al., 2023). To prepare *N. parisii* spores, *C. elegans* N2 worms were infected with *N. parisii* on NGM plates and incubated for several days to generate a large infected population. Infected worms were harvested, confirmed to not contain contaminating bacteria, and stored at −80°C. To purify spores, worms were mechanically disrupted with 2 mm zirconia beads and the homogenate was passed through a 5 μm filter (Millipore). Purified spore preparations were confirmed to be free of bacterial and fungal contamination and stored at −80°C, and spore concentrations were determined by counting DY96-stained spores on a Cell-VU counting slide.

### Infection assays

Continuous infection assays were carried out in 24-well plates in a total volume of 400 μL containing 800 L1 worms, 1% DMSO, and 12,000 *N. parisii* spores/μL. Each compound was tested at 10 μM, 40 μM, and 100 μM. Plates were sealed with breathable adhesive film, wrapped in parafilm, and incubated at 21°C with shaking at 180 rpm for 4 days. Worms were then washed in M9 containing 0.1% Tween-20, fixed in acetone, and stained with DY96, which binds the chitin of microsporidia spores and *C. elegans* embryos. Stained samples were imaged by fluorescence microscopy on a ZEISS Axio Imager 2, and images were captured using Zeiss Zen 2.3. Animals were scored as gravid if they contained any embryos and as infected if they displayed any newly formed spores. Severe infection was defined as spores observed >50% of the worm’s intestine. Smaller L1 progeny of the treated animals were excluded from all measurements.

### Statistical analysis

Data were collected from three independent biological replicates and analyzed in R version 4.5.2 and RStudio 2026.1.1.403 (R Development Core Team, 2019). For each compound, infection or gravidity was compared to the corresponding DMSO control at each concentration by one-way ANOVA followed by Dunnett’s post-hoc test. Normality of the data was assessed for each dataset using ANOVA residuals. Because the residuals deviated from normality, for all but the data in Fig. 2B, a non-parametric Kruskal-Wallis test with Dunn’s post-hoc comparison to the control was also performed, and yielded consistent conclusions for the major effects. Results of all statistical tests are included in Table S1.

## 3. Results

Small-molecule inhibitors are a classical means of manipulating protein function in living organisms (Stockwell, 2004). To explore the susceptibility of microsporidia to such compounds, we identified 15 inhibitors that have known protein targets in *Saccharomyces cerevisiae* and have not been previously tested against *N. parisii* (Ottilie et al., 2022; Smith et al., 2010). We also included three inhibitors—fumagillin, albendazole, and dexrazoxane—that have previously been shown to have activity against *N. parisii* (Huang et al., 2023; Murareanu et al., 2022).

To test the activity of these inhibitors against *N. parisii*, we used a 24-well liquid infection assay in which synchronized first larval stage (L1) worms were cultured with *N. parisii* spores in the presence of each compound at 10, 40, or 100 μM. After fixation and staining with Direct Yellow 96 (DY96), we quantified both the percentage of animals that were infected and the percentage displaying severe infection. Consistent with their previously reported activity, albendazole, fumagillin, and dexrazoxane significantly reduced infection at multiple concentrations, decreasing infection to below 10% at 100 μM (Fig. 1A). Most other compounds did not significantly reduce infection, although tunicamycin strongly inhibited infection and camptothecin, etoposide, and geldanamycin produced weaker, but significant effects (Fig. 1A). When we restricted our analysis to only severely infected worms at the 100 μM drug dose, 15 of the 18 inhibitors had significant effects, however, only albendazole, fumagillin, dexrazoxane, and tunicamycin strongly reduced infection (Fig. 1B) (Huang et al., 2026a).

**Fig. 1.**
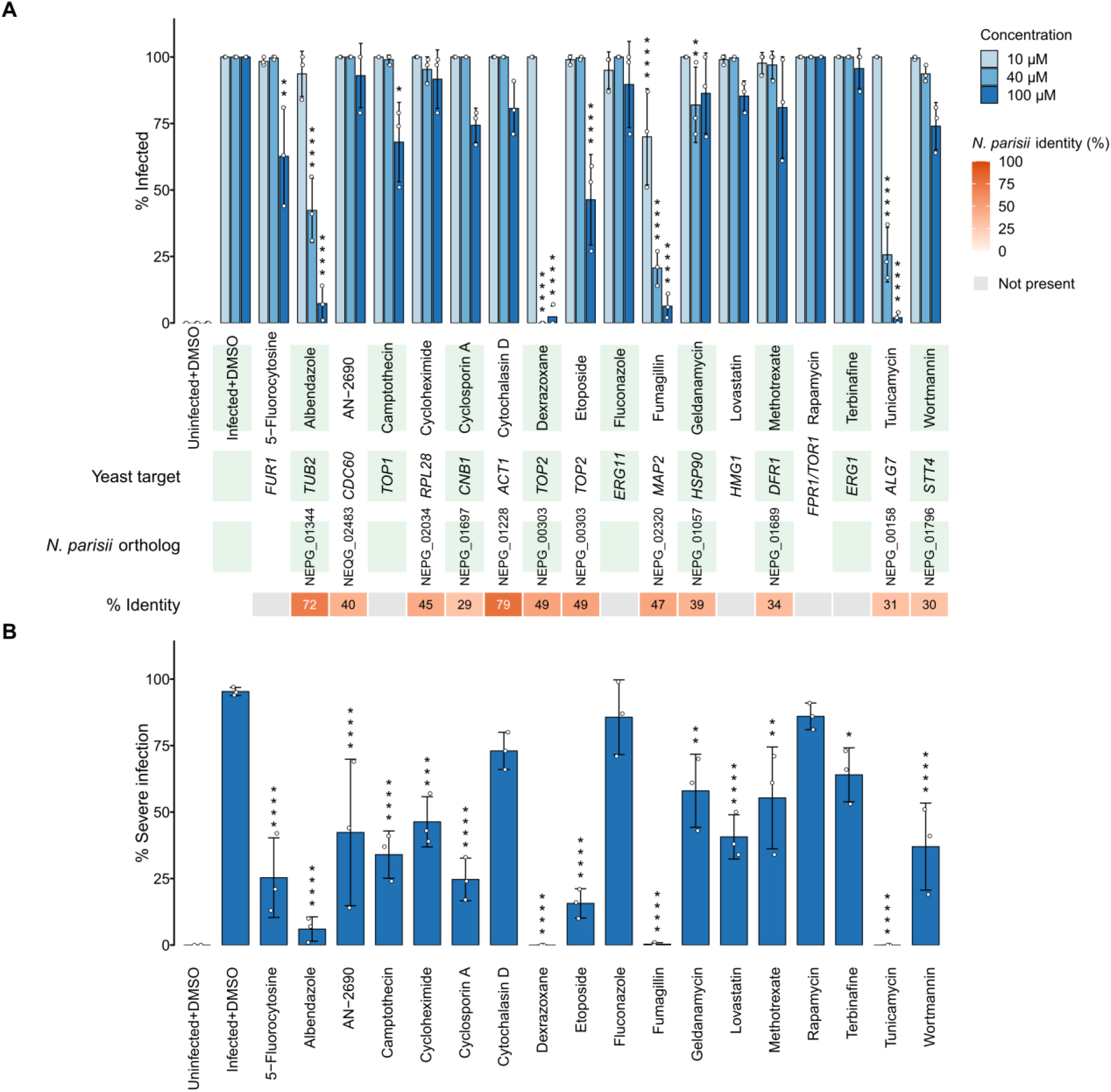
Effect of compounds on *N. parisii* infection of *C. elegans*. **(A)** Percentage of infected animals following treatment with the indicated compounds at 10, 40, and 100 µM (light to dark blue). Below the graph, the yeast protein target, the corresponding *N. parisii* ortholog, and its percent amino acid identity to the yeast protein are indicated; “Not present” denotes compounds with no identifiable *N. parisii* ortholog. Identification of microsporidia orthologs and conservation identity was done either using https://blast.ncbi.nlm.nih.gov/Blast.cgi or taken from Jiang et al. (Jiang et al., 2025). **(B)** Percentage of animals with severe infection (spores present throughout >50% of the worm’s intestine) following treatment at 100 µM. In both panels, bars show the mean of three biological replicates, individual replicates are shown as points, and error bars represent the standard deviation (SD). n = 3 biological replicates, N = ≥ 100 worms counted per biological replicate. Statistical significance was assessed by one-way ANOVA followed by Dunnett’s post-hoc test comparing each treatment to the infected DMSO control at the corresponding concentration (*, P < 0.05; **, P < 0.01; ***, P < 0.001; ****, P < 0.0001).

Infection of *C. elegans* with *N. parisii* reduces reproductive fitness, and inhibitors can rescue this phenotype (Murareanu et al., 2022). To assess rescue, we used gravidity—the presence of embryos—as a metric of reproductive fitness. Significant rescue of gravidity at multiple concentrations was observed only for albendazole, dexrazoxane, and fumagillin, with weaker but significant rescue at 100 μM for wortmannin (Fig. 2A). Tunicamycin produced no rescue despite inhibiting infection, so we tested all inhibitors in uninfected animals across the three concentrations. Tunicamycin eliminated gravidity at every concentration, indicating overt host toxicity (Fig. 2B).

**Fig. 2.**
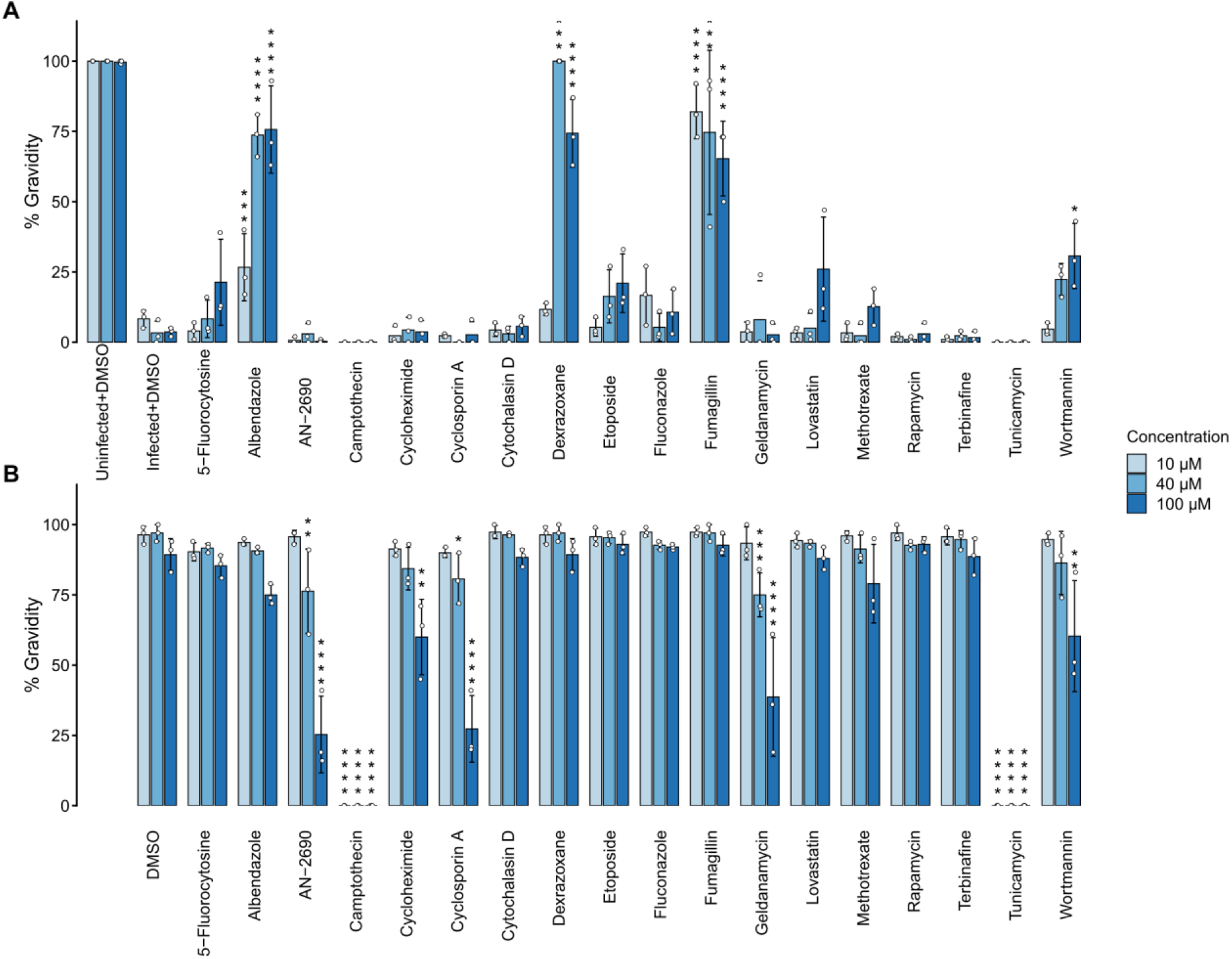
Effect of compounds on *C. elegans* gravidity. **(A)** Percentage of gravid animals after exposure to *N. parisii* and **(B)** percentage of gravid uninfected animals following treatment with the indicated compounds at 10, 40, and 100 µM (light to dark blue). In both panels, bars show the mean of three biological replicates, individual replicates are shown as points, and error bars represent the standard deviation (SD). n = 3 biological replicates, N = ≥ 100 worms counted per biological replicate. Statistical significance was assessed by one-way ANOVA followed by Dunnett’s post-hoc test comparing each treatment to the corresponding DMSO control at the same concentration (Infected+DMSO for panel A, DMSO for panel B; *, P < 0.05; **, P < 0.01; ***, P < 0.001; ****, P < 0.0001).

Comparing inhibitor efficacy with target conservation revealed no simple relationship between the two (Fig. 1A). The most effective compounds spanned a wide range of target conservation: albendazole, dexrazoxane, fumagillin, and tunicamycin all act on yeast targets which have identifiable *N. parisii* orthologs, but with widely varying identities (beta-tubulin [TUB2], 72%; topoisomerase II [TOP2], 49%; methionine aminopeptidase 2 [MAP2], 47%; UDP-N-acetylglucosamine-1-phosphate transferase [ALG7], 31%). Many other inhibitors with targets in this same range of conservation produced little or no inhibition. Conversely, several compounds whose targets have no *N. parisii* ortholog still weakly reduced infection.

## 4. Discussion

We assembled a panel of 18 small-molecule inhibitors with characterized protein targets in *S. cerevisiae* and tested their ability to inhibit *N. parisii* infection of *C. elegans*. Of the 15 compounds not previously tested in this system, only tunicamycin strongly inhibited infection. Most of these compounds have not previously been reported to be tested against any microsporidian species; the few exceptions are cytochalasin D, shown to inhibit *Encephalitozoon hellem* spore germination, cyclosporin A, shown not to significantly inhibit *Encephalitozoon cuniculi*, and tunicamycin, shown to inhibit *Encephalitozoon intestinalis* growth (Didier et al., 2013; Lallo et al., 2013; Leitch et al., 1993). Almost all inhibitors reduced severe infection to some degree, but only at the highest concentration tested and ranging from 45–85% inhibition; none were as effective as the previously characterized compounds albendazole, fumagillin, and dexrazoxane, and none strongly restored the reproductive fitness of *C. elegans*. The lack of a clear correlation between target conservation and inhibitory activity suggests that sequence identity is a poor predictor of whether a compound will inhibit *N. parisii*, and that factors such as the conservation of specific drug-contacting residues and compound uptake likely play a larger role in determining efficacy. Consistent with the latter, *C. elegans* has a robust capacity to detoxify small molecules, which renders some compounds ineffective in this animal (Roy, 2025).

Tunicamycin inhibits the first step of N-linked glycosylation by targeting ALG7 in *S. cerevisiae* and is a classical pharmacological inducer of endoplasmic reticulum stress and the unfolded protein response (UPR) (Shamu et al., 1994). The *N. parisii* ALG7 ortholog is only 31% identical to its *S. cerevisiae* counterpart, and ALG7 from *Trachipleistophora hominis* or the microsporidian relative *Rozella allomycis* could not complement an *S. cerevisiae* ALG7 deletion strain (Jiang et al., 2025). Tunicamycin has previously been shown to cause developmental delay, and we likewise observed a potent reduction in embryo production (Denzel et al., 2014). Although tunicamycin is likely inhibiting microsporidia directly, further experiments will be needed to confirm this and to determine whether inhibition of the host UPR affects *N. parisii* growth. Little is known about endoplasmic reticulum stress in microsporidia, and this compound could be a useful tool for investigating it.

## Acknowledgements

We thank Gio Dela Cruz and P. M. Shreenidhi for helpful comments on this manuscript. This work was supported by Canadian Institutes of Health Research grant (no. 461807 to A. W. R.) and Q. H. was supported by an award from the China Scholarship Council.

## Data availability

All data are in Data S1.

## Declaration of generative AI and AI-assisted technologies in the manuscript preparation process

During the preparation of this work the authors used Claude Opus 4.8 and ChatGPT 5.5 to generate R scripts for figures and tables, to generate draft text for the paper, and to provide a comprehensive literature review. The authors reviewed and edited the content and take full responsibility for the content of the published article.

## Author contributions

Q. H. and A. W. R. conceptualized the experiments. Q. H. performed all the experiments. Q.H. and A. W. R. analyzed the experimental data. The paper was written by A. W. R., with edits from all authors. Mentorship and funding acquisition were provided by A. W. R.

